# W2RAP: a pipeline for high quality, robust assemblies of large complex genomes from short read data

**DOI:** 10.1101/110999

**Authors:** Bernardo J. Clavijo, Gonzalo Garcia Accinelli, Jonathan Wright, Darren Heavens, Katie Barr, Luis Yanes, Federica Di-Palma

## Abstract

Producing high-quality whole-genome shotgun *de novo* assemblies from plant and animal species with large and complex genomes using low-cost short read sequencing technologies remains a challenge. But when the right sequencing data, with appropriate quality control, is assembled using approaches focused on robustness of the process rather than maximization of a single metric such as the usual contiguity estimators, good quality assemblies with informative value for comparative analyses can be produced. Here we present a complete method described from data generation and qc all the way up to scaffold of complex genomes using Illumina short reads and its application to data from plants and human datasets. We show how to use the w2rap pipeline following a metric-guided approach to produce cost-effective assemblies. The assemblies are highly accurate, provide good coverage of the genome and show good short range contiguity. Our pipeline has already enabled the rapid, cost-effective generation of *de novo* genome assemblies from large, polyploid crop species with a focus on comparative genomics.

**Availability:** w2rap is available under MIT license, with some subcomponents under GPL-licenses. A ready-to-run docker with all software pre-requisites and example data is also available.

http://github.com/bioinfologics/w2rap

http://github.com/bioinfologics/w2rap-contigger

## 1 Introduction

Generation of a high quality genome assembly is a crucial first-step towards understanding the biology of an organism. It establishes a complete catalogue of genes and provides the foundation for characterising the genetic variation within a species and how this variation impacts gene function and phenotypic variation. Over the last 10 years, many methods have been described to address this problem and the genomes of many organisms have been published. Yet, for plant and animal species which often have large and complex genomes, assembly remains a fundamental challenge.

Genome assemblies generated from massively parallel short-read technologies such as Illumina are highly accurate at the nucleotide level and relatively inexpensive to generate, but remain highly fragmented due to complex repeat content and varying degrees of polymorphism and ploidy. While solutions such as ALLPATHS-LG (Gnerre et al., 2010), have revolutionised the field enabling the sequencing and assembly of many mammalian genomes and providing a foundation for large scale comparative analysis and lineage-specific evolutionary analysis, they require a precise recipe of input libraries coupled to a fixed set of algorithm parameters which are not suitable for larger, complex genomes.

Here we present a pipeline called w2rap (Wheat/Whole-genome Robust Assembly Pipeline) to rapidly generate high-quality, low-cost, robust assemblies from genomes with different levels of complexity. Our approach uses Illumina PCR-free paired end (PE) 250bp reads for contig construction with the w2rap-contigger, an improved algorithm based on DISCOVAR *denovo* (Weisenfeld et al., 2014) (Love et al., 2016), and Nextera long mate-pair (LMP) libraries for long-range scaffolding with SOAPdenovo2 (Luo et al., 2012). W2rap encompasses a full data processing workflow from raw reads to scaffolds, and crucially allows the user to fine tune the algorithmic parameters making draft assembly generation an iterative process adaptable to diverse genome complexities and data. We also show in our w2rap test *A. thaliana* dataset that tuning assembly to enhance accuracy produces more contiguous assemblies at lower computational cost, as the downstream analysis problems become easier.

We demonstrate our approach here by applying it to a datasets from *Ara-bidopsys thaliana* and show how it performs in line with state of the art approaches in standard *Homo sapiens* data. We have already used used it to assemble the hexaploid, highly repetitive, 17Gbp *Triticum aestivum* (bread wheat) genome which generated highly accurate scaffolds in agreement with the existing single chromosome reference sequence (Clavijo et al., 2016). Our results maintain the completeness and accuracy achieved by DISCOVAR *denovo* coupled with reduced memory usage, and processing time. Most importantly, increased accuracy and contiguity are achieved by enhanced parameterisation of the algorithms, improved repeat resolution, and the systematic use of LMP data via SOAPdenovo2 scaffolding. This method makes it possible to consistently generate high-quality draft assemblies for large, complex genomes at low cost.

## 2 Results

### 2.1 Data generation

W2rap uses a combination of Illumina Paired End (PE) and Long Mate Pair (LMP) reads. We recommend PCR-free PE libraries, using fragment sizes of about 700bp for optimal short-repeat resolution in the w2rap-contigger stage. Nextera LMP libraries (Heavens et al., 2015) are used to provide precise and cost-effective longer range information.

Whilst different combinations of coverages and library sizes can be used depending on genome complexity, repeat structure, and other characteristics, we recommend a minimum coverage of 30x PE and up to 100x for highly heterozygous genomes. For LMP libraries, a minimum coverage of 30x raw reads per library is recommended, with up to 50x being routinely used in internal projects. For plant genomes that often contain high levels of LTR-retrotransposons, it is important to include a LMP library longer than 8Kbp to span these highly repetitive blocks. We have successfully used a combination of libraries of 8Kbp, 10Kbp and 14Kbp for complex genomes such as wheat. As a general rule, when trying to iteratively improve an assembly by sequencing a range of LMP libraries, we recommend getting 30x LMP coverage of a library with an insert size corresponding to the N80 of the current scaffolding results.

### 2.2 Assembly quality and contiguity

When all algorithmic heuristics perform as intended, the genome assembly results should become more accurate, and this accuracy can drive contiguity. Table 1 and Figure 1 show a clear example of an *A. thaliana* assembly becoming more accurate and contiguous, by changing parameters of the same heuristics to achieve better processing. The scaffolding results from both the *A. thaliana* contigs and scaffolds outperform both ABySS and SOAP for our test runs on contiguity, and outperform DISCOVAR *denovo* contigs for both accuracy and contiguity. Details of the assembly parameters for each of these runs are given on Supplementary Material Section 5.

**Table 1:** Comparison between w2rap and other assembly tools for the A. *thaliana* dataset. All values in Kbp.

| Assembler | Contig NG50 | Contig NGA50 | Scaffold NG50 | Scaffold NGA50 |
| --- | --- | --- | --- | --- |
| ABYSS v1.9 | 57 | 56.57 | 412.58 | 370.42 |
| SOAPdenovo2 | 21.5 | 21.49 | 937 | 365.86 |
| w2rap | 361.9 | 355.89 | 1318.81 | 1136.96 |

**Figure 1:**
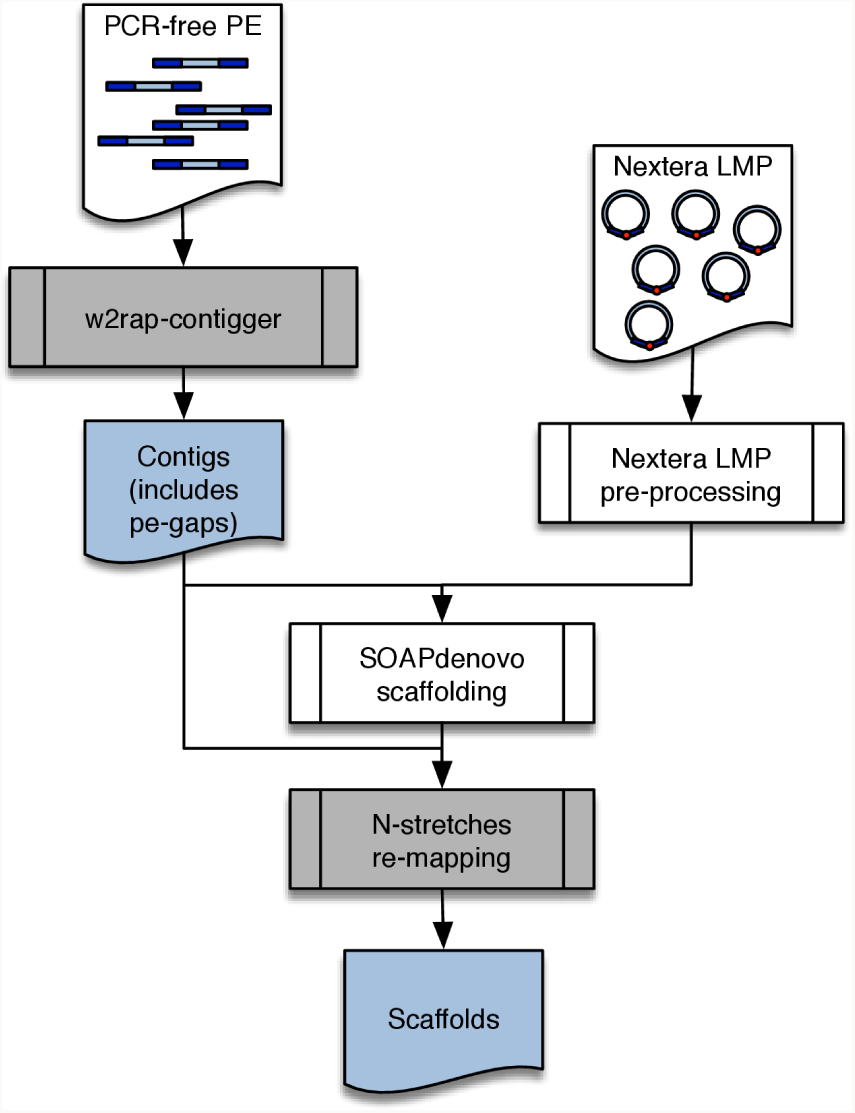
w2rap assembly workflow. Contigs are produced by running the w2rap-contigger on PCR-free 2x250bp Illumina data. Nextera Long Mate Paired reads are then pre-processed with the w2rap scripts and used to scaffold the contigs using the SOAPdenovo2 scaffolder included in w2rap. A script then re-introduces N-runs from the original contigs displaced during SOAPdenovo2 scaffolding.

We also used the human HG004 Illumina 2x250bp PE and LMP datasets from the Genome in a Bottle Consortium (Zook et al., 2016), and we cite for comparison results from the ABySS 2.0 publication (Jackman et al., 2016). It is important to highlight the main focus of w2rap is producing assemblies for comparative studies. Therefore, it is crucial to produce assemblies that are consistently comparable and not one-off optimal settings for a particular dataset. The default w2rap-contigger parameters achieve similar results to those of DISCOVAR *denovo*, as they run essentially the same heuristics with near-identical parameter choice, but achieve slightly less contiguity as seen for the HG004 dataset in Table 2. This comes from slightly more conservative choices of parameters. A ‘dv_like’ mode can be activated when running the w2rap-contigger that produces results more similar to those of DISCOVAR *denovo*. The general results from these assemblies are comparable to those mentioned on the ABySS 2.0 publication.

**Table 2:** Comparison between w2rap with other assembly tools for the HG004 *H. sapiens* dataset from Genome in a Bottle.

| Assembler | Contig NG50 (Kbp) | Scaffold NG50 (Mbp) |
| --- | --- | --- |
| ABYSS v1.9 | 30.0 | 4.36 |
| SOAPdenovo2 | 3.8 | 0.17 |
| w2rap | 90.6 | 4.78 |

\* Data for ABYSS and SOAPdenovo2 results from ABYSS 2.0 preprint.

### 2.3 Computational performance of w2rap-contigger

Among the disadvantages of using DISCOVAR *denovo* to assemble large and complex genomes are its high memory consumption and long runtime and for projects aiming to generate multiple assemblies computational resources can become a major bottleneck. We reviewed the whole codebase and re-implemented many algorithms with openMP-based parallel approaches, testing and improving the performance on NUMA systems, such as those needed to handle the memory and processing requirements of larger genome assemblies. This led to reduced processing times, both in NUMA systems and normal servers. Further reduction on the processing times can be achieved by correct parametrisation of the first steps of the contig assembly process, as this leads to simpler problems on the later stages (See Supplementary Material for more details).

Each step during contig assembly uses significantly different algorithmic approaches and data. We segmented the w2rap-contigger processing into eight steps which can be run independently thus enabling us to make more efficient usage of resources when running multiple assemblies or sharing computational resources with other projects. This change produced two desired outcomes: (i) each step runs with the resources required for that step only, thus avoiding a waste of computing resources on large-memory multi-processor machines and, (ii) the granularity of running shorter steps rather than all steps combined allows for better control over the assembly, and provides the opportunity for a detailed check of results from intermediate steps. These modifications are important when assembling large and complex genomes, where the contigging steps can take over 10 days.

## 3 Methods

### 3.1 Data generation

PCR-free Illumina paired end libraries are both cost-effective and less prone to representation bias (Huptas et al., 2016). Standard Illumina NGS library construction protocols target fragments with a 500bp insert size but these typically span 300bp to 700bp, and smaller molecules within a library are more likely to be sequenced over larger molecules. These libraries reduce effective coverage due to overlapping sequences and fail to provide the spatial information due to the predominance short insert fragments. For *de novo* genome assembly projects, greater accuracy and contiguity can be achieved if the unique sequence flanking repeats can be resolved within multiple single paired reads. We aim for fragments of 700bp and longer which can be achieved by using a more stringent AMPure XP bead based clean up (for reproducibility it is recommended to use a positive displacement pipette to accurately dispense the beads) or a size exclusion technology such as the Blue Pippin from Sage Science.

We recommend following our modifications to the Nextera LMP protocol to produce libraries with good size distributions and representation (Heavens et al., 2015). Processing of these is explained in a later section, but emphasis must be on QC. When sequencing large genomes, we recommend constructing multiple libraries, sequencing a combined pool in a single low-coverage (and low cost) run, then choose the libraries with the best characteristics for further sequencing.

### 3.2 Generating contigs with the w2rap-contigger

The w2rap-contigger is a extensively modified version of DISCOVAR *denovo*. The original DISCOVAR has some limitations in terms of large repeat-rich datasets, which made it impossible to run it on genomes such as that of hexaploid wheat. We fixed bugs and limitations on the repeat-resolution heuristics and implemented an extra repeat-resolution heuristic, the PathFinder, that is described in the Supplementary Material. We divided the original assembly heuristics into discrete steps, both to optimise the usage of computational resources and to make the heuristic processing easier to track. Supplementary Material Section 1 contains more detail about each step.

The w2rap-contigger provides a more extensive set of parameters than that of DISCOVAR *denovo,* although most of these were originally present in the code but fixed to values that were reasonable for mammalian genomes as sequenced by The Broad Institute. In general, when sequencing multiple related genomes for comparative studies, a sequencing recipe should be devised and then a set of parameters chosen for the whole study, to guarantee comparable results.

While trying to adjust parameters for contiguity is a widespread practice in genome assembly, we have shown in the results section and Figure 1 that increasing accuracy can lead to higher contiguity. This means the assembly process must be guided by the careful execution of each heuristic to achieve accuracy. There needs to be an understanding of each heuristic and a method to measure whether each heuristic is achieving the desired results.

#### 3.2.1 Understanding the w2rap-contigger metrics

At the beginning and end of each step, and at every relevant point during execution, the w2rap-contigger prints a set of assembly status metrics: kmers in the graph, graph contiguity, reads pathing (i.e. a single-end read placement) and pair status. (See Supplementary Material for details).

The w2rap-contigger represents the assembly graph internally as a list of edges (sequences or gaps) with a list of vertices representing K-1 overlaps. The graph is directed, with independent reverse complements, which means every sequence will be represented both in forward and reverse unless it is palindromic. This means the number of kmers in the graph will be roughly doubled. Alongside the number of kmers and edges, a set of Nk20, Nk50 and Nk80 values show the length in kmers of the edges such that edges of that length or more cover 20%, 50% and 80% of the total kmer size of the graph. We use this value rather than the traditional NXX values because it makes sense to evaluate edge lengths using a kmer-based method.

In terms of read paths and pairs status, as the assembly progresses we should generally see an improvement in the number of ends that map to a unique location and the number of pairs with both ends mapped and satisfied. These are the more indicative metrics, and we recommend following them throughout all the steps described below. A more detailed and up-to-date explanation on how to use the metrics to guide the assembly will always be kept in the w2rap tutorial.

#### 3.2.2 60-mer graph construction

The first three steps on the w2rap-contigger transform the reads into its binary format (step 1), produce a 60-mer count with neighbouring information (step 2) and construct a 60-mer graph (step 3). The main parameter to adjust in these steps is the minimum frequency of 60-mers. This can be adjusted on step 2, and then can be re-adjusted to higher values on step 3 if needed on high coverage datasets where errors are over-abundant. The 60-mer spectrum is written to the small_K.freqs file, and we recommend choosing values of minimum frequency smaller than the first valley, which separates the bulk of the error distribution from the bulk of the genome's true 60-mers. At this step it is also advisable to check the fragment size distribution, saved in small_K.frag.dist and control that small improvements on NkXX are not achieved by loosing placement for too many reads.

#### 3.2.3 Optimising large_K value

At this point, a collection of all possible paths through the graph are generated, effectively transforming the 60-mer graph into an exploded large-K graph that contains all possible large kmer paths. This graph is then evaluated for support, pruning paths that do not have support from the original reads. The main parameter for steps 4 and 5 is the kmer size for the second (and final) graph. This value again represents a trade-off between connectivity and noise. Increasing this parameter will disentangle repeats in the kmer spectrum, but will also generate more erroneous paths, thus decreasing the amount of reads mapped to the true paths and eventually making them discontiguous due to lack of support. A secondary parameter, representing the number of supporting reads required for a path to be considered valid is also available.

#### 3.2.4 Optimising local assembly

At this point, a key requirement is to maximise the graph connectivity so the desired assembly can be found as a path through the graph. As previous steps may have incorrectly pruned true paths, a local heuristic to reconstruct them is now used. First, clusters of unsatisfied read pairs are generated: these are reads that connect a set of “left” and a set of “right” edges in the graph, which are not currently connected through existing paths. All left edges need to be connected between them and all right edges needs to be connected between them. The set of these ’’bridging” reads is assembled using the local assembly methods described in the DISCOVAR *denovo* publication. A selection of possible paths through the previously unrepresented region is created. A new sequence graph is computed by including all the edges from the previous graph and the edges from the local assembly heuristics. This graph is the basis of the final assembly.

The number of reads in each cluster is sub-sampled to a fixed value. This is reasonable both because it places an upper-bound on computational resources, and because the reads are only spanning a region between two ends of a pair, which means a region of less than 1Kbp. The number of read pairs is defaulted to 200, but this can be adjusted with the pair_sample parameter. While increasing this parameter may in some cases improve the assembly, it is computationally expensive.

#### 3.2.5 Optimising repeat resolution

This step cleans all artifactual paths and edges in the graph and attempts to resolve repeats by using read-mapping information through the edges. There are 2 different heuristics for repeat resolution which can be used in different combinations and with different parameters;

- PullAparter: inherited from the original DISCOVAR *denovo* heuristics, expands edges with 2 neighbours on each side, where there is read evidence to separate the two instances of the repeat. This method has been optimised to run faster on complex genomes but the simple heuristics remain the same.
- PathFinder: expands loops using a combination of read support and coverage-based heuristics, then looks for single-flow repeat regions. These are regions where a number of small, complex connected paths, flow in a single direction from a set of N inputs into a set of N outputs, and where all these inputs and outputs are assumed to be unique sequence on the assembly. Reads mapping from the input edges to the output edges is then evaluated to score in-out combinations. If an in-out 1-to-1 combination with correct support is found, and it is possible to reconstruct the path through the single-flow region based on read mappings, then the whole region is expanded and solved for each 1-to-1 pairing (see Supplementary figure).

The dv_like mode of the contigger runs just the PullAparter, like the original DISCOVAR. The default mode runs the PullAparter (to solve the easier cases) and then the PathFinder.

#### 3.2.6 Parameterisation

It is important to note that modifying parameters at each stage of the w2rap-contigger may significantly affect runtime. If the first 60-mer graph produces a clean, highly resolved assembly, many of the complex heuristics for cleanup and specially for local assembly and repeat resolution won't need to be used to analyse the bulk of the graph. This again highlights the value of being able to run the algorithms in a consistent manner with metrics that show when the parameters are being set correctly.

### 3.3 Scaffolding

#### 3.3.1 Preparing Nextera LMP reads

The pipeline for processing Nextera LMP reads is based on Nextclip (Leggett et al., 2013) and is designed to recover correctly generated LMP reads from raw sequencing reads. The Nexera protocol generates circularised constructs containing the Nextera adapter at the junction between the two reads. However, the adapter doesn't necessarily occur at the end of each read. To account for this, reads are first combined into a single sequence where they overlap using FLASh (Magoc and Salzberg, 2011) to generate a single sequence which should contain the Nextera adapter. This sequence is then reverse complemented to generate the other read in the pair. Nextclip (Leggett et al., 2013) is used to classify reads according to whether the adapter is found and where in the read-pair it is located. This whole process is encapsulated within a single script provided as part of the pipeline.

The K-mer Analysis Toolkit can be used to compare the LMP reads to the PE reads to highlight sequence representation issues. A subset of the reads are then mapped to the previously assembled contigs to check the fragment size distribution.

#### 3.3.2 Optimising LMP mapping and scaffolding

A bundled version of SOAPdenovo is used for scaffolding. The main parameters at this stage are the kmer size used to map the reads and the read support to call a link, as in the original SOAP scaffolder. A kmer size of 71 is reasonable in most cases of complex genomes, but checking read placement stats and link status on the output files is recommended. Finally, s_scaff is used to generate scaffolds. The output file generated at this stage gives useful information about how the scaffolding performed.

#### 3.3.3 Recovering gaps and creating releases

Before scaffolding, SOAPdenovo converts gaps in contigs (Ns) to Cs and Gs so these are converted back to Ns by mapping the contigs back to scaffolds using the output files from SOAPdenovo. A script is provided for this as outlined in the tutorial.

When deciding on a length cut-off for scaffolds in a final release, the K-mer Analysis Toolkit can be used to make sure this doesn't result in a loss of content. The scaffolds should also be checked for contamination (such as phiX) and Illumina adapters.

**Figure 2:**
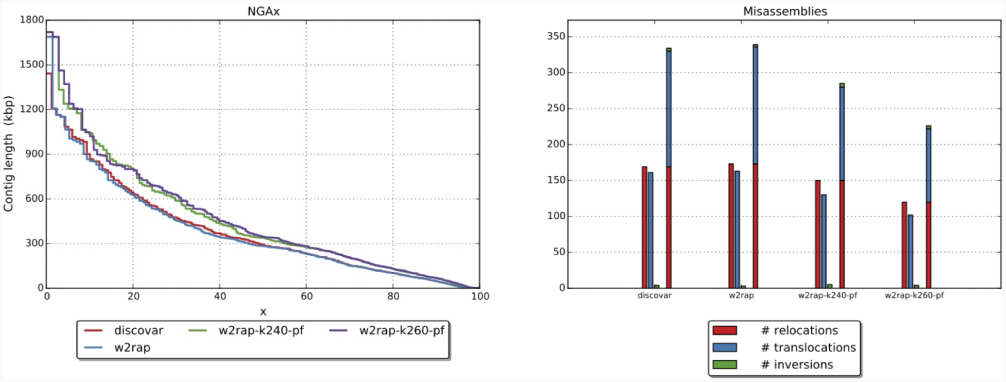
Correct parametrisation improves contiguity and accuracy. Contigs generated using K=260 and the PathFinder heuristic show increased aligned contiguity when evaluated against the TAIR10 reference. At the same time the correct parametrisation shows a decrease in misassemblies, which indicates improved performance from the algorithms instead of just contiguity gains by joining less-supported links.

## 4 Conclusion

Our assembly method to construct contigs and scaffolds from short read Illumina data produces high-quality assemblies in a cost-effective way, unlocking information in complex genomes. The focus on metrics enables complete tracking of how the datasets and algorithms are performing, which becomes particularly important for comparative studies where multiple similar genomes are assembled from equivalent datasets. By using w2rap, performance of the whole assembly process can be tracked to ensure reproducibility.

We have shown the effectiveness of combining the w2rap-contigger's short-range accuracy, based on the DISCOVAR heuristics originally designed to preserve variation on the assembly datasets, with a quality focused long mate paired sequencing method and the simple but proven heuristics of SOAPdenovo2’s scaffolding modules. While more expensive or specific approaches could produce particular one-off results outmatching w2rap’s performance, these assemblies are a good starting point for many comparative genomics projects where robustness, accuracy and price are the most important factors to consider. As shown by its performance on the *H. sapiens* dataset, w2rap also scales well with complexity, and is already in use for even more complex genomes including the highly complex, 17Gbp genome of hexaploid wheat.

## 5 Author contributions

BJC designed the assembly approach and the pipeline. BJC and GGA programmed the w2rap-contigger. JW, GGA and BJC programmed the scripts for the w2rap pipeline. DH tweaked sequencing protocols and produced test datasets. BJC, LY and GGA tuned and optimised software, compilation chains and architecture. BJC, GGA, JW and KB tested assembly results and evaluated the pipeline. FDP provided initial collaboration with the Broad Institute and feedback throughout the project. BJC and FDP wrote the manuscript, with contributions from all authors.

## 6 Acknowledgments

Thanks to David Jaffe and Neil Weisenfeld for their support on reusing the DISCOVAR codebase. Thanks to the EI PP team for continuous efforts on producing great data, and many helpful discussions and feedback. Thanks to all members of the BBSRC Wheat LOLA for continuous feedback and support.

This work was strategically funded by the BBSRC, Institute Strategic Programme Grant BB/J004669/1. Work on wheat assembly was funded by BBSRC strategic LOLA Award BB/J003743/1. This research was supported in part by the NBIP Computing infrastructure for Science (CiS) group.

